# Exercise alleviates liver senescence but does not outmatch the effect of dietary restriction in diet-induced MASLD

**DOI:** 10.1101/2024.12.03.626591

**Authors:** Angeliki Katsarou, Grigorios Papadopoulos, Ioannis I. Moustakas, Argyro Papadopetraki, Athanasios Moustogiannis, Aigli-Ioanna Legaki, Eirini Giannousi, Dimitris Veroutis, Athanassios Kotsinas, Vassilis G Gorgoulis, Anastassios Philippou, Michael Koutsilieris, Antonios Chatzigeorgiou

**Affiliations:** Department of Physiology, Medical School, National and Kapodistrian University of Athens, 11527 Athens, Greece; Molecular Carcinogenesis Group, Department of Histology and Embryology, Medical School, National and Kapodistrian University of Athens, 11527 Athens, Greece; Department of Histology and Embryology, Medical School, National and Kapodistrian University of Athens, 11527 Athens, Greece; Ninewells Hospital and Medical School, University of Dundee, Dundee, UK; Molecular and Clinical Cancer Sciences, Manchester Cancer Research Centre, Manchester Academic Health Sciences Centre, University of Manchester, Manchester, UK; Biomedical Research Foundation, Academy of Athens, Athens, Greece

## Abstract

**Background:** The present study aims at deciphering the potential benefits of aerobic exercise and dietary restriction on liver senescence, which is an established hallmark of metabolic dysfunction-associated steatotic liver disease (MASLD), a condition with limited therapeutic options.

**Methods:** C57BL6 mice were subjected to normal diet (ND, 10% of kilocalories from fat) or a high-fat diet (HFD, 60% of kilocalories deriving from fat and water supplemented with 5% High-fructose Corn Syrup, HFCS) for 12 weeks. Then, for additional 8 weeks, the ND group continued with the same diet, while the HFD group was divided into four subgroups: a) mice that continued with the same HFD-5% HFCS in water scheme (HFD), b) mice that continued with the same HFD-5% HFCS in water scheme and underwent supervised aerobic exercise 3-times/week (HFDEX), c) mice that were switched to ND (dietary restriction, DR) and d) mice that were switched to ND while undergoing supervised aerobic exercise 3-times/week (DREX). Phenotypic and histological characterization of obesity and MASLD were performed in all groups. Biomarkers of senescence were analyzed in terms of their mRNA expression levels to assess the impact of all interventions on MASLD-related senescence in the liver. GL13 and p21 immunohistochemical stainings were conducted to examine the protein levels of senescence-associated lipofuscin and p21^WAF1/CIP1^ respectively, so as to finally investigate their relationship with the grade of steatosis observed in each individual animal.

**Results:** DR and DREX groups exhibited significantly reduced features of obesity and MASLD-related hepatic steatosis, to a greater extent than the respective amelioration driven by aerobic exercise-only in HFDEX animals. A statistically significant increase of the mRNA expression of cyclin-dependent kinase p21^WAF1/CIP1^ was detected in HFD livers as compared to ND, which was also reversed upon DR-inclusive interventions. In contrast, the gene expression levels of cyclin-dependent kinase p16^INK4a^ remained similar in all groups even after a combined intervention. Increased hepatic expression of the p27 and p53 components of the p53-p21 ^CIP/WAF^-driven axis of cellular senescence as well as their restoration to ND-like levels upon DR and DREX, suggest an active participation of the p21^WAF1/CIP1^ mechanism of senescence in the emergence of MASLD, but also in its reversal through DR or/and EX interventions. Immunohistochemical stainings _for_ GL13 and p21 confirmed the aforementioned alterations of p21^WAF1/CIP1^ at the tissular level.

**Conclusion:** Liver senescence is responsive both to exercise and dietary restriction, but its amelioration in the context of MASLD is more robust upon DR-inclusive interventions.

## 1. Introduction

Senescence represents a cellular state characterized by the inability for further proliferation or growth[1]. During senescence, cells retain an active metabolic role, usually related to the senescence-associated secretory phenotype (SASP). Specifically, SASP, a phenotype acquired by cells undergoing irreversible cell cycle arrest, might introduce dynamic changes in several cellular processes happening within the senescent cell’s adjacent microenvironment, through secretion of cytokines, chemokines, growth factors and even extracellular matrix components[1]. In addition, senescent cells overexpress cell cycle inhibitors such as p16^INK4a^, p21^WAF1/CIP1^and p53, which are main drivers of cycle arrest at phase G1 and ultimately, can drive accumulation of unrepaired DNA damage[2]. Senescence may result from aging and/or chronic stress-related stimuli[1, 2]. Therefore, senescence retains major physiological importance in metabolic organs, such as the liver, where pathology involves fluctuations in several traditional components of SASP, like pro-inflammatory cytokines and extracellular matrix mediators (e.g. matrix metalloproteinases, MMPs)[1, 2]. In the case of metabolic dysfunction-associated steatotic liver disease (MASLD), a chronic liver disease with broad phenotypic spectrum, stress-induced hepatic senescence can occur due to lipid accumulation, subsequent peroxidation of lipid species and aberrant production of reactive oxygen species[1, 2]. Importantly, senescence has also been proposed as a prerequisite for progression of liver steatosis during aging, in murine models[3]. Interestingly, liver senescence has been also correlated with the severity of fibrosis, as observed in human liver biopsies, and often molecular hallmarks of senescence such as p21^WAF1/CIP1^, have been shown to predict disease progression[3]. In the present study, we investigated the potential nature of liver senescence as manifested in diet-induced MASLD but mostly, beneath the prism of its potential ‘alignment’ with the recession of MASLD through dietary and exercise-inclusive interventions.

## 2. Materials and Methods

Information related to Materials and Methods utilized in this study are found in the Supporting Information.

## 3. Results

### 3.1 Hepatic steatosis is ameliorated upon exercise or/and dietary fat restriction during MASLD

In order to investigate the independent or combined effect of exercise and dietary fat restriction on liver senescence, this study included five groups of C57BL/6J mice as illustrated in **Fig.1a**. Specifically, The first group followed a normal diet (ND) with 10% of kilocalories originating from fat and the second a high-fat diet (60% of kilocalories from fat), supplemented with 5% high fructose corn syrup (HFCS) in water (defined as a high-fat diet group, HFD) for 20 weeks. The HFD group which was initially fed a high-fat diet for a 12-week period, was afterwards split into four distinct groups; one group continued the same feeding conditions, while the rest three followed a different intervention for the remaining 8 weeks of the experiment. Specifically, one sub-group was subjected to an 8-week intervention of three weekly aerobic exercise sessions (HFDEX), one switched from a high-fat to a normal diet (dietary restriction, DR), and the final group underwent a combination of dietary restriction and the same aforementioned aerobic exercise regimen (DREX). The dietary restriction intervention was utilized as a simplistic reduction of dietary fat consumption.

**Fig 1.**
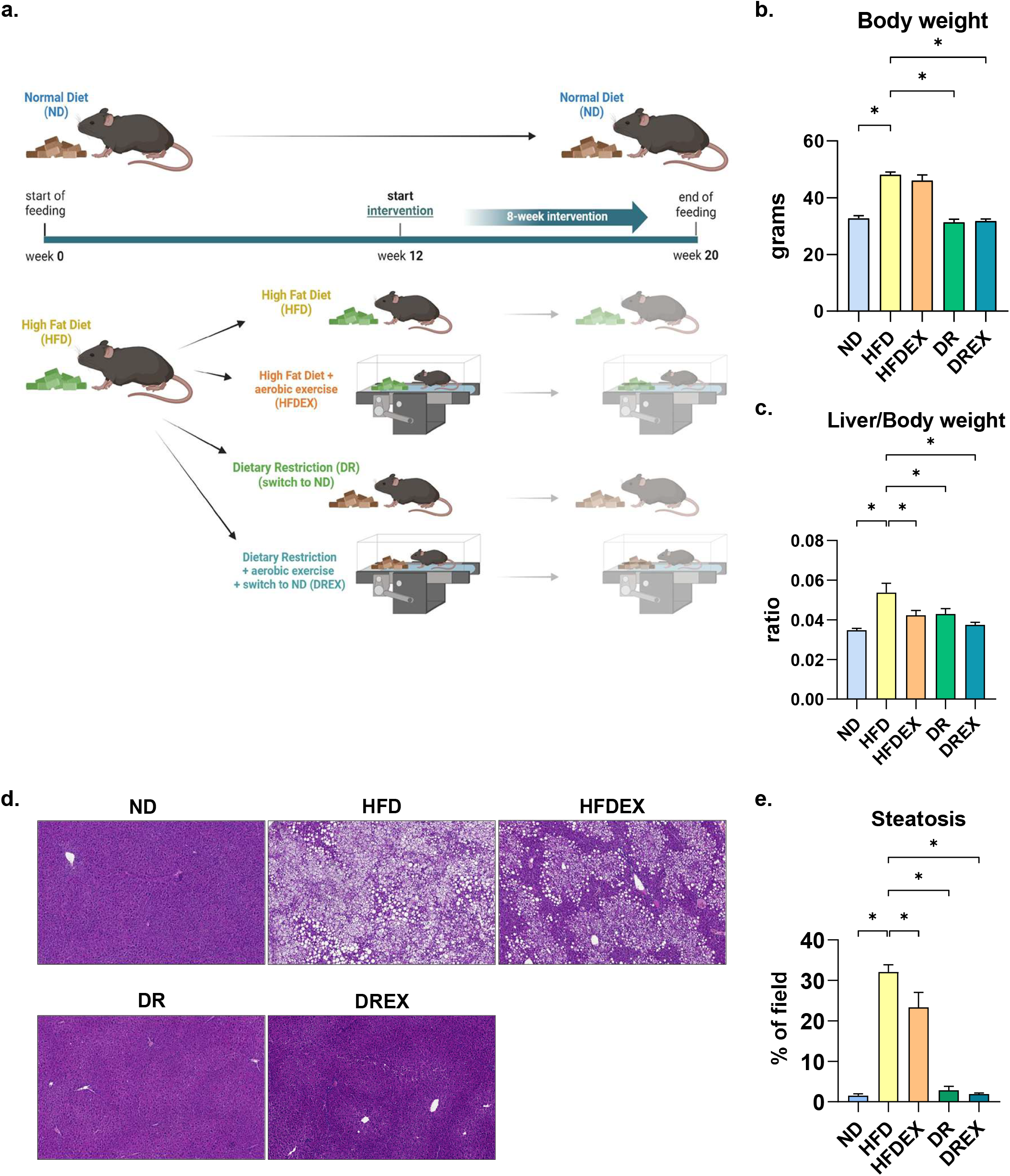
Dietary Restriction and Aerobic Exercise mitigate hepatic steatosis during MASLD. **a**. Experimental outline of the interventions’ protocol. **b**. Body weights of mice in distinct experimental groups at the end of the feeding protocol. **c**. Liver-to-body weight ratios of mice in distinct groups at the end of the feeding protocol. **d**. Representative images from hematoxylin and eosin (H&E)-stained liver tissue sections (magnification at 100x). **e**. Hepatic steatosis levels in the distinct groups expressed as percentage of the lipid droplet area per field of vision derived from H&E-stained images. *n= 6-10 mice/group. Data are mean ± SEM. *: p < 0*.*05*.

At the end of the experiment, the induction of obesity, measured as body weight increase in HFD mice, was significantly reversed in the DR intervention groups, while HFDEX mice exhibited a sensibly milder reduction in their body weight (**Fig.1b**). Following animal sacrification, the livers of mice were excised and weighed, revealing a significant reduction in the liver-to-body weight ratio across all intervention groups (**Fig.1c**). Hematoxylin-Eosin (H&E) staining of liver tissue sections was used to quantify steatosis as percentage of lipid droplets per tissue field (**Fig.1d**), confirming that the severe levels of steatosis in HFD, exhibited statistically significant reduction upon DR, DREX, and even upon EX alone, albeit with a less robust effect in the latter (**Fig.1e**). This implies that the individual intervention of aerobic exercise might not be optimal for managing obesity, but is sufficient to ameliorate severe steatosis even when applied independently.

### 3.2 Dietary restriction is superior to exercise for ameliorating liver senescence during MASLD

To examine the potential of DR and EX to counteract liver senescence, a main hallmark of MASLD, the expression of several senescence-related genes was studied in each group. Strikingly, each one of the DR and DREX interventions resulted in significantly reduced p21^WAF1/CIP1^ expression in the livers of the respective mice, as compared to the expression levels in HFD mice (**Fig.2a**). A similar, but not statistically significant trend of reduction was observed regarding the expression levels of p53 in all three interventions as compared to HFD livers (**Fig.2a**). However, the respective mRNA expression levels of p16^INK4a^ did not show any substantial differences among the groups (**Fig.2a**), pinpointing that the senescent compartment of the liver in the presented phenomenon might rely more on the p53-p21^WAF1/CIP1^ axis of stress-induced senescence. Regarding hepatic expression levels of p27, a marker of both senescence and regeneration in the liver, was observed to be significantly reduced solely upon combination of DR and EX (**Fig.2a**).

**Fig 2.**
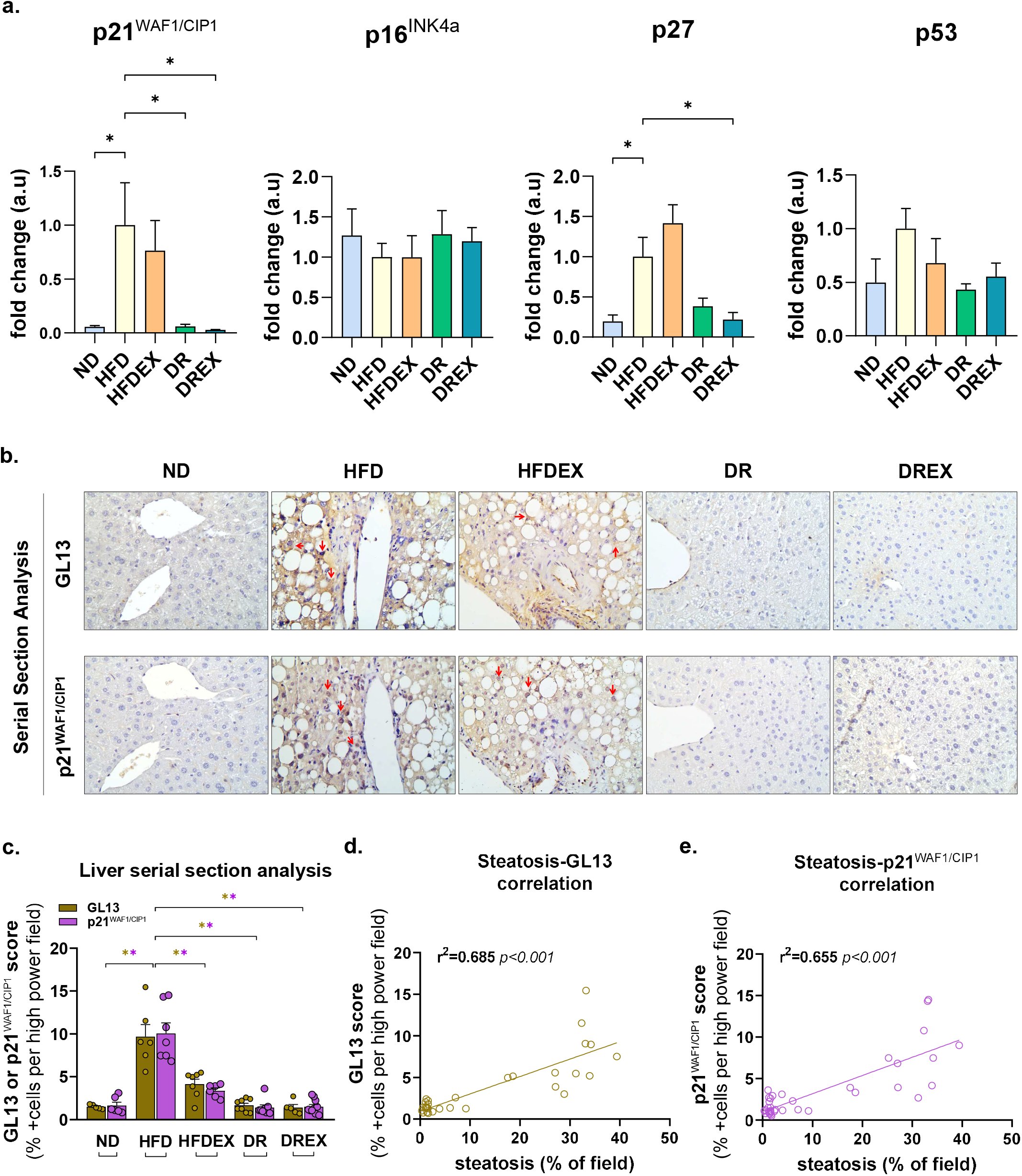
MASLD-associated liver senescence is responsive to both dietary and aerobic exercise interventions with dietary restriction being superior. **a**. mRNA expression levels of p21^WAF1/CIP1^, p16^INK4a^, p27 and p53 in liver tissues of the five groups assessed with qPCR and quantified by using eukaryotic translation elongation factor 2 (EEF2) gene as an internal control. Expression levels were normalized to the HFD group, which was set as 1. **b**. Representative immunohistochemistry images of the senescent markers GL13 and p21^WAF1/CIP1^ in serial sections from mouse liver tissues, counterstained with hematoxylin. Red arrows indicate the presence of lipofuscin, denoted by GL13 staining (strong brown cytoplasmic signal), or the nuclear localization of p21^WAF1/CIP1^(strong brown cytoplasmic signal), in separate serial sections. **c**. Average GL13 and p21 ^WAF1/CIP1^ scores per experimental group expressed as percentages of positively stained cells per high-power field. **d-e**. Pearson’s correlation of GL13 and p21^WAF1/CIP1^ scores with liver steatosis levels of individual mice from all experimental groups. *n= 4-10 mice/group, Data are mean ± SEM. *: p < 0*.*05*.

In turn, the presence of senescence was studied immunohistochemically in the liver tissues. GL13/p21^WAF1/CIP1^ tissue-positive immunostaining was used to detect lipofuscin-rich cells that contain nuclear p21^WAF1/CIP1^; an estimation of the percentage of +cells per high-power field was performed in serial liver sections (**Fig.2b**). The percentage of GL13-positive cells – indicative of the presence of lipofuscin in the liver tissue – was higher in the HFD mice, but its presence was diminished in livers of all intervention groups (**Fig.2c**). Importantly, a high degree of positive correlation of hepatic steatosis grade with both GL13 and p21^WAF1/CIP1^ was also evident (**Fig.2c-e**), thus, underpinning a potential mechanistic role of p21^WAF1/CIP1^ -driven senescence within the context of the studied interventions and their anti-MASLD beneficial effects.

## 4. Discussion

Herein, we demonstrate that obesity-related MASLD provoked by a HFD/HFCS feeding in mice is related to a p21^WAF1/CIP1^ -associated senescent hepatic phenotype, which can be partially or rigorously attenuated through lifestyle interventions including dietary fat restriction, exercise, or a combination of both. Optimal results pertinent to amelioration of hepatic senescence, were achieved upon DR alone or in combination with exercise, while exercise-alone yielded a less pronounced effect. Along this line, DR provoked notable changes in senescent-related markers such as p21^WAF1/CIP1^ and p27; though, p16^INK4a^ did not follow the same pattern among the groups, implying that the p21^WAF1/CIP1^ axis of cellular senescence might be orchestrating the presented phenomenon.

Nowadays, human studies have highlighted the beneficial role of weight loss, diet composition, and physical activity in the management of MASLD; thus, we attempted to simulate these first-line interventions that are commonly utilized for MASLD management[4]. Data from murine studies support that even low-intensity exercise is beneficial for liver health; specifically, low and moderate treadmill exercise in mice could ameliorate liver fibrosis[5]. However, the mechanisms *via* which these non-pharmaceutical interventions influence different stages of MASLD remain unclear, particularly those connecting exercise to MASLD prevention. In addition, data on the effect of simultaneous dietary and exercise regimens are also scarce.

Evidence supports the benefits of the above interventions in reducing liver steatosis, in parallel to the amelioration of hepatic senescence. The diet composition seems to play a primary role, since a diet with high protein intake and fat can exaggerate the presence of hepatic senescence[6]. A study utilizing murine models fed a fast-food diet showed that exercise led to a significant reduction of hepatic p21^WAF1/CIP1^ expression; yet, this pattern was not observed for p16^INK4a^ and p53[7]. Our team has previously reported hepatic senescence involvement in MASLD development regardless of the presence of obesity by proving the presence of senescence both in obese and non-obese NAFLD models[8].

A more recent study in NF-kB1^-/-^ mice that are prone to aging and liver disease showed that modest aerobic exercise ameliorated hepatic steatosis; simultaneously, the intervention led to the decline of p21^WAF1/CIP1^-positive cells in liver sections and hepatocytes with damaged telomeres, compared to the group subjected to normal activity[9]. Similarly, another study in Sprague-Dawley rats found that the combination of exercise and caloric restriction was proven superior at reducing liver lipid accumulation, when compared to exercise or diet alone[10]. Additionally, caloric restriction was more efficient than exercise but with no statistical significance; nevertheless, all interventions significantly reduced hepatic steatosis compared to the HFD group, similar to the findings of the present study[10]. The senescence-associated β-gal activity was limited compared to the increased activity in the HFD[10]. In the same direction, the expression of p16^INK4a^ and p27 in liver tissue was suppressed with exercise and diet, while the combination of both emerged superior[10].

Our study proposes a significant benefit of common lifestyle interventions against MASLD-related senescence. This is demonstrated by the reduced expression of p21^WAF1/CIP1^ and p27, and secondarily of p53 which did not reach significance, markers that are robustly expressed during obese-MASLD, but not of p16^INK4a^, upon dietary restriction-inclusive interventions. Exercise alone also significantly restored the hepatic p21^WAF1/CIP1^ expression towards the levels of the healthy group, but with a milder decrease, as our immunohistochemical analysis showed. The combinatory intervention (DREX) induced similar significant changes in p21^WAF1/CIP1^ and p27. Consequently, this could imply that the mechanism of MASLD amelioration might involve regulation of the p53-p21^WAF1/CIP1^ axis, rather than the p16^INK4a^ pathway. However, it should be reminded that senescence is likely a prerequisite in both ageing- and diet-induced MASLD and can manifest with high heterogeneity within different phenotypes.

## Acknowledgements

The work was supported by grants from the Hellenic Foundation for Research & Innovation (HFRI) and the Hellenic Association for the Study of the Liver (HASL), both to A.C.

