## Supplementary Materials and Methods for "Exercise alleviates liver senescence but does not outmatch the effect of dietary restriction in diet-induced MASLD"

**Katsarou, Papadopoulos et al.**

**Materials and Methods**

**Animal studies**

Mice were housed at 20-22 oC on a 12h light/dark cycle with ad libitum access to food and water. Male C57BL/6 mice (obtained from the Hellenic Pasteur Institute, Athens, Greece) at 8-10 weeks of age were fed either a normal diet (ND) with 10% of kilocalories from fat and tap water, or a high-fat diet (HFD) containing 60% of kilocalories from fat (D12450B and D12492, respectively, Research Diets, New Brunswick, NJ, USA), while the latter received water supplemented with 5% High fructose Corn Syrup 55 (HFCS) for induction of obesity-related MASLD [1] (Best Flavors, CA, USA), for 12 weeks. After 12 weeks, mice previously subjected to HFD and 5% HFCS were divided into four subgroups: a) mice which continued with the same HFD plus 5% HFCS in water (HFD), b) mice which continued with the same HFD plus 5% HFCS in water and underwent sessions of supervised aerobic exercise three times /week (HFDEX), c) mice which were switched to ND and tap water (dietary fat restriction, DR) and d) mice which were switched to ND and tap water and were concurrently put into three sessions of supervised aerobic exercise weekly (DREX). All four subgroups, as well as the initial ND group continued feeding as described above, for an additional 8 weeks. Of note, the HFDEX and DREX feeding groups underwent three 30-minute aerobic exercise sessions on a weekly basis on a PanLab LE8700 Treadmill. Mice were urged to achieve a running speed of 20 cm/sec. At the end of the feeding period, blood was collected and the mice were euthanized and subjected to systemic perfusion with PBS, and livers were excised and weighed. The animal work was approved by the Region of Attica, Greece.

**Histopathology**

Freshly isolated liver pieces were fixed using 10% formalin solution and embedded in paraffin. Then, 5-µm-thick sections were prepared and were subjected to hematoxylin/eosin (H&E) staining. A computerized Olympus Slideview VS200 slide scanner was utilized for picture acquisition. For assessing steatosis, the percentage of lipid droplets per field of vision was evaluated in the H&E-stained liver sections, by using Fiji ImageJ software [2].

**RNA Isolation and qPCR**

Total RNA from livers was extracted using TRItidy G (APPLICHEM, A0451). After quantification of RNA concentration, cDNA synthesis was performed by utilizing the PrimeScript™ RT Reagent Kit (Takara, Shiga, Japan, RR037). The PowerUp Sybr Green™ Master Mix (Thermo Fischer Scientific, Applied Biosystems) was used to conduct qPCR on a QuantStudio™ 5 Real Time PCR System (Thermo Fischer Scientific, Applied Biosystems). The relative mRNA expression was calculated according to the ∆∆Ct method [3] by using the expression of eukaryotic translation elongation factor 2 (EEF2) gene as an internal control.

**Immunohistochemistry (IHC)**

FFPE tissues were cut in 4μm sections and stained for detection of senescent cells using GL13 reagent (SenTraGor) and the detection of p21^WAF1/Cip1^ protein as described previously [4]. Briefly, sections were deparaffinized, hydrated and antigen retrieval was heat-mediated using citric acid buffer (pH=6) for 15 minutes in steamer for GL13 staining or 25 minutes in microwave in the case of p21^WAF1/Cip1^. Samples were cooled for 20 minutes on ice bath. Normal goat serum was used (abcam, ab138478) in a 1:40 dilution for blocking non-specific binding sites of antibodies. Endogenous biotin was blocked using the Biotin blocking kit (SP-2001) according to manufacturer’s instructions. GL13 was applied for 10 minutes at 37^o^C and primary anti-biotin antibody (ab201341, abcam) was applied in dilution 1:300 at 4^o^C overnight. For p21^WAF1/Cip1^ staining after cooling step tissue section was treated with the primary anti-p21^WAF1/Cip1^ (ab188224, abcam) in dilution 1:200 at 4^o^C overnight. Then positive signal in all samples was developed using the Dako REAL EnVision Detection System, (Cat.no: K5007) according to the manufacturer’s instructions. Specimens were counterstained with Hematoxylin. The whole segments were scanned and the mean percentage of GL13 or p21^WAF1/Cip1^ positive cells was obtained from 5-10 high power fields [5].

**Statistical Analysis**

For all comparisons, One-Way Analysis of Variance (ANOVA) with Tukey’s correction was performed. Pearson’s correlation was utilized for examining the concomitant presence of GL13 or of the p21^WAF1/CIP1^ protein with the grade of hepatic steatosis per individual mouse. In the case of qPCR experiments, outliers were identified and removed within the distinct experimental groups using Grubb’s test. GraphPad Prism v8.0.1 software was used. Data are expressed as mean ± SEM and p-value was set as p < 0.05.

**Τέλος φόρμας**
